# Heterotrophic diazotrophy along a river–lake continuum: lifestyle and contribution to N_2_ fixation

**DOI:** 10.1101/2024.09.25.614945

**Authors:** Eyal Geisler, Hagar Siebner, Max Kolton, Guy Sisma-Ventura, Eyal Rahav, Shai Arnon, Edo Bar-Zeev

## Abstract

Heterotrophic diazotrophs are potentially important agents in freshwater ecosystems, yet they remain poorly understood. This study elucidates the contribution of freshwater heterotrophic diazotrophs as free-living or aggregate-associated cells to total N_2_ fixation along the continuum from the Jordan River to Lake Kinneret, Israel. Heterotrophic diazotrophs accounted for 25%–56% of the total diazotrophs and commonly found as free-living cells or attached to aggregates in the river. N_2_ fixation by heterotrophic diazotrophs associated with aggregates varied along the river, while accounting for ~50% of N_2_ fixation in the lake. Non-cyanobacterial diazotrophs dominated the free-living fraction in the river, and cyanobacteria were mostly identified in the lake. The diversity of free-living diazotrophs was sensitive to environmental changes, as the aggregates have likely provided a protected micro-environment. The results reveal the dynamic lifestyle of heterotrophic diazotrophs along a river–lake continuum and highlight their contribution to total N_2_ fixation and primary production.

**Teaser:** Freshwater heterotrophic diazotrophs are more ubiquitous than previously thought, can be found as free-living cells or associated with aggregates, and significantly contribute to primary production.

## Introduction

Bioconversion of dissolved dinitrogen (N_2_) into biologically available forms is a fundamental biochemical process in aquatic environments and often controls ecosystem primary productivity (*1*). Biological N_2_ fixation is catalyzed by the enzyme nitrogenase, a metalloprotein complex found in a specific group of bacterial and archaeal prokaryotes known as diazotrophs (*2*). These microbes inhabit a wide range of aquatic environments, including the open ocean (*3*), coastal areas (*4*), estuaries (*5*), rivers (*6*), lakes (*7*), and man-made reservoirs (*8*). In contrast to the open ocean, the physicochemical structure of freshwater systems is highly heterogeneous and strongly influenced by anthropogenic inputs and climate change (*9*).

### BOX.

*specific definitions*

**Heterotrophic diazotrophs:** Bacteria and archaea that use organic matter from external sources to sustain their metabolic requirements for N_2_ fixation.

**Non-cyanobacterial diazotrophs (NCDs):** The term NCDs has been widely used in the literature, yet these microbes are only phylogenetically classified as cyanobacteria and non-cyanobacteria. From a metabolic perspective, NCDs are highly diverse and occur as photoheterotrophic, heterotrophic, chemolithoautotrophic, and/or mixotrophic diazotrophs.

Most knowledge about aquatic diazotrophs has come from oceans because the consensus over recent decades has been that N_2_ fixation rates in freshwater environments are low, and thus do not have a substantial effect on N inputs (*10*). However, because freshwater and terrestrial environments are closely linked, even low rates of N_2_ fixation can alter ecological stoichiometry and microbial activity, thereby affecting primary production (PP) and secondary production (*10*). Regardless of the type of aquatic ecosystem, diazotroph activity is limited by several key factors (*1*), including a lack of energy-rich molecules (e.g., ATP and NADPH) for nitrogenases to reduce N_2_ to ammonia; elevated concentrations of dissolved oxygen, which impair the nitrogenase catalytic site; the limited availability of nutrients (e.g., iron and phosphorus) and trace elements (e.g., molybdenum); and excessive concentrations of dissolved inorganic nitrogen (DIN). These factors often dictate the governing lifestyle (e.g., free-living or aggregate-associated), activity, diversity, and metabolic pathways within an aquatic environment.

Aquatic studies have mostly focused on cyanobacterial diazotrophs (*11*), which meet their energy requirements via photosynthetic carbon fixation and use various strategies to protect nitrogenase (*1*). These phototrophic diazotrophs are often captured as planktonic bloom-forming filamentous chains (*12*) and unicellular cells (*13*) as well as dense mats (*14*) attached to different types of benthic substrates, such as rock and wood (*15*). Although sparsely studied three decades ago (*16*), the possible importance of planktonic heterotrophic diazotrophs in freshwater N_2_ fixation has only recently been reported by some research groups (*14*, *17*, *18*). These freshwater heterotrophic diazotrophs <u>were</u> mostly affiliated with clusters I\III and often include *Gammaproteobacteria spp.* and *Thermodesulfobacteriota spp.* The *nifH* clusters represent evolutionary lineages of diazotrophs divided by aerobic and anaerobic metabolisms and characterization level (*19*). Despite the potential importance of diazotrophs to freshwater ecosystems, few studies have measured their N_2_ fixation rates, undertaken diazotroph identification, quantified *nif*H genes copies (*10*). Moreover, there has been little investigation of the environmental factors that govern the lifestyle of heterotrophic diazotrophs along the river–lake continuum.

In this study, we determined the contribution of free-living and aggregate-associated heterotrophic diazotrophs to total N_2_ fixation along the Jordan River-to-Lake Kinneret continuum. An approach coupling flow-cytometry with nitrogenase immunolabeling was developed to quantify the number of diazotrophs. N_2_ fixation rates were determined by enriching water samples with dissolved ^15^N_2_ while simultaneously analyzing diazotroph diversity using *nifH* amplicons. The results show the effects of various abiotic drivers on diazotrophy and highlight the major contribution of free-living and aggregate-associated heterotrophic diazotrophs to N_2_ fixation in this freshwater environment.

## Materials and methods

### Sample collection and preparation

Surface water was collected at sites along the Jordan River, denoted as the upstream (Up), midstream (Mid), and downstream (Down) sites, and from the margin of Lake Kinneret, denoted as the lake site (Figure 1). In 2020–2021, two sampling campaigns were conducted in summer and two in winter. Hydrological data, such as river flow and water level, were retrieved from the Israeli Water Authority (https://floods.online). Surface water temperature and turbidity were measured *in situ* using a sensor (WTW, Multi 3420, Xylem Analytics, Washington, DC, USA). Additional parameters were measured directly at each site, including the concentrations of large particles (>150 µm), total organic carbon (TOC), total nitrogen (TN), inorganic nutrients (NH_4_^+^, NO_2_^−^ + NO_3_^−^, and PO_4_^3−^), chlorophyll α (Chl *a*), and transparent exopolymer particles (TEP).

**Figure 1.**
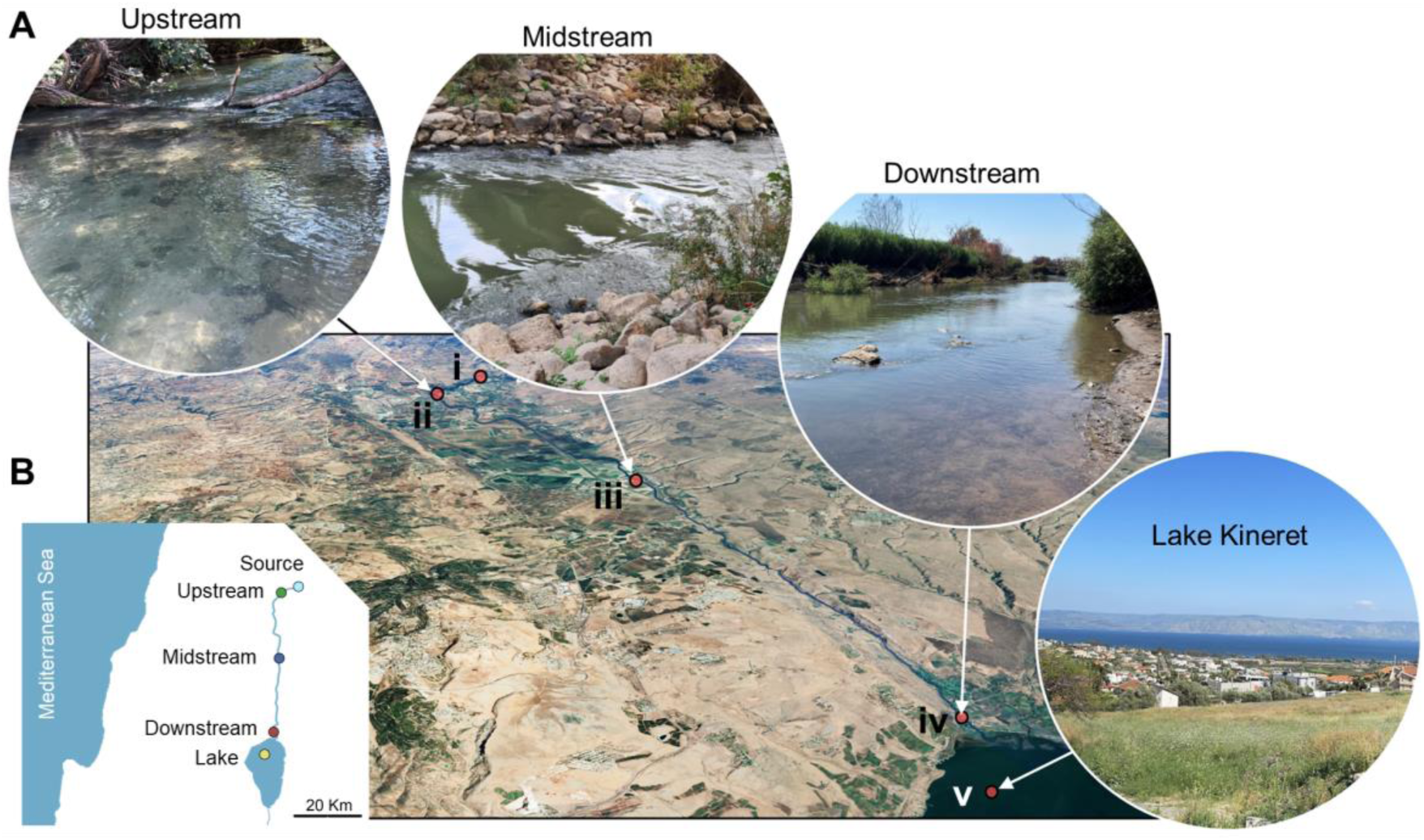
(A) Oblique satellite image showing the sampling sites along the Jordan River-to-Lake Kinneret continuum and (B) schematic map of the research area. The perennial source of the Jordan River is the Tel-Dan Spring (i). The upstream sampling site (ii) is located 2.6 km from Tel-Dan Spring, one of the main sources of the Jordan River. The midstream site (iii) is located downstream of a 25-km section of river that passes through an area of intensive agriculture. The downstream site (iv) is located 16 km downriver of the midstream site, close to where the river discharges into Lake Kinneret (the overall length of the Jordan River is ~44 km). Surface water from Lake Kinneret was collected 1–2 km from the shore (v).

Surface water from each site was sampled in five 1 L transparent Nalgene bottles. Four bottles were supplemented with enriched dissolved ^15^N_2_ water, and one bottle was filled with sample water only to determine the natural abundance of ^15^N_2_. Three of the four bottles supplemented with ^15^N_2_ also had 50 µM of –[3,4-dichlorophenyl]-1,1-dimethylurea (DCMU; D2425, Sigma-Aldrich, St. Louis, MO, USA) added as a photosynthetic inhibitor and were incubated under dark conditions to analyze heterotrophic diazotrophy (*4*). The fourth bottle was incubated under ambient light without a photosynthetic inhibitor to estimate both phototrophic and heterotrophic diazotrophy. All bottles were incubated under ambient temperature with gentle shaking (G10, New Brunswick Scientific Co) at ~3 xg. for 48 h. At the end of the incubation period, the water samples were filtered gently using a volume pump (<300 mbar) and a 12 µm cellulose filter (WHA10400012, Whatman, Merck, Darmstadt, Germany). The collected material (particles of ≥12 µm), consisting of aggregates and their associated microbiome, was resuspended in sterile pre-filtered ambient water (100 mL, 0.2 µm filter). The filtrate, containing particles and cells smaller than 12 µm, remained in the original 1 L Nalgene bottles. Analyses for both the aggregate-associated (≥12 µm) and free-living (filtrate, <12 µm) fractions included bacterial production (BP), bacterial abundance (BA), diazotrophic abundance (DA), and N_2_ fixation rates.

### Analytical approaches

#### TEP concentration

Water samples (25–100 mL, four technical replicates for each site) were filtered onto a polycarbonate filter (0.4 µm, GVS, Life Sciences, Bologna, Italy) until the meniscus and was dyed with 5% Alcian blue stain (Sigma-Aldrich) and stored at −20 °C in glass scintillation vials within a week. TEP were extracted by soaking the filters in 80% sulfuric acid (5 mL, Carlo Erba, Cornaredo, Italy) for 2 h, after which the extracted solution was measured spectrophotometrically at 787 nm. A five-point calibration curve with gum xanthan stock (GX; Sigma G1253, Sigma-Aldrich) was used to estimate TEP concentrations (*20*).

#### Concentrations of nitrogen species and orthophosphate

Water samples were pre-filtered using 0.4 µm polycarbonate filters and kept at −20 °C in acid-washed plastic vials until analysis. Orthophosphate (PO_4_^3−^), NO_3_^−^ + NO_2_^−^, and ammonium (NH_4_^+^) were measured using a segmented flow autoanalyzer system (AA-3, Seal Analytical, Wrexham, UK) equipped with colorimetric detection. The nutrient limit of detection, defined as three times the standard deviation of 10 measurements of the blank, were 8, 80, and 90 nM for PO_4_^3−^, NO_3_^−^ + NO_2_^−^, and NH_4_^+^, respectively. The reproducibility of the analyses was determined by using the following certified reference materials (CRM): MOOS 3 (PO_4_^3−^, and NO_3_^−^ + NO_2_^−^), VKI 4.1 (NO_3_^−^ + NO_2_^−^), and VKI 4.2 (PO_4_^3−^). Results were used when measured CRMs were within ±10% of the certified values. All concentrations were well above their respective detection limit (*21*).

#### TOC and TN concentrations

Water samples were collected in acidified 40 mL glass tubes (1 M hydrochloric acid, pH = 2.5) and stored at −20 °C until analysis with a TOC and TN analyzer (Multi N/C, Analytik Jena, Jena, Germany; detection limit = 0.3 mg L^−1^). Before each measurement, a calibration curve was constructed using TOC (1090170100, Merck) and NH_4_^+^ (1198120500, Merck) standards, ranging from 0 to 100 mg L^−1^. Organic nitrogen (N_org_) was calculated by subtracting the sum of inorganic nitrogen species (NO_2_^−^, NO_3_^−^, and NH_4_^+^) from the TN.

#### Particle size and concentration

Freshly collected water samples (200 mL) were analyzed in triplicate for particle concentrations in units of particles per liter (>150–2200 µm) by using a particle counter (WGS-267 Water Grab Sampler, Met One, Grants Pass, OR, USA) at a flow rate of 100 mL min^−1^.

#### Chl a concentration

A sample (60 mL) was filtered onto glass fiber filters (GF/F; 1825-025 Lifegene, Mevo Horon, Israel) and the filters were stored at −20 °C under dark conditions until analysis. Chl *a* was extracted overnight in the dark with 90% acetone solution and was then measured by the non-acidification method as described in Welschmeyer, 1994 using a fluorometer (Trilogy, Turner Designs, San Jose, CA, USA) equipped with 436 nm excitation and 680 nm emission filters. Chl *a* standard from *Anacystis nidulas* (Sigma C6144, Sigma-Aldrich) was used to calibrate the fluorometer. Blank filters treated the same way were also tested and showed readings below the detection limit (<0.01 µg L^−1^) (*22*).

#### BP analysis

Triplicate samples (1.7 mL) were spiked with a mixture of radiolabeled:unlabeled leucine solution. The leucine solution contained 30 nM ^3^H leucine (NET1166001MC, PerkinElmer, Waltham, MA, USA) and 70 nM of non-radioactive leucine (Sigma L-8000, Sigma-Aldrich), making a total of 100 nmol leucine L^−1^ per sample. The samples were incubated in the dark at ambient temperature for 4 h before being halted with trichloroacetic acid (TCA; 100 µL) solution (100%). Killed blanks containing sample water, leucine mixture, and TCA without incubation were also taken at each site, and their reads were subtracted from the non-killed reads. In the lab, the samples were analyzed using the micro-centrifugation technique following Smith & Azam (1992) and mixed with Ultima Gold scintillation cocktail (L8286, Sigma-Aldrich) and (1 mL). Radioactive counts per minute were measured with a scintillation counter (Tri-Carb 4810 TR, <u>PerkinElmer</u>) and converted into carbon using a factor of 1.5 kg C mol^−1^ and an isotope dilution factor of 2 (*23*).

#### PP analysis

Triplicate samples (50 mL) were spiked with 5 µCi NaH^14^CO_3_ (NEC086H005MC, PerkinElmer) and incubated for 6 h under dark or ambient light conditions, representing chemoautotrophy and photosynthesis, respectively. River/lake water spiked with NaH^14^CO_3_ without incubation was also analyzed, and the readings were subtracted from the sample reads. The samples were filtered onto GF/F filters and the filters were left with 50 µl of 32 % of hydrochloric acid overnight to remove inorganic carbon residues. The filters were mixed with Ultima Gold scintillation cocktail (5 mL) analyzed with the scintillation counter as described above (*24*).

#### N_2_ fixation rates

Samples (1 L) were incubated with ^15^N_2_-enriched freshwater (7.5% *v:v*) for 48 h under dark (including DCMU) or light conditions. The enriched ^15^N_2_ water was prepared and kept in a 1 L transparent Nalgene bottle with a gas-tight cap until use within a few days (see the Supporting Information for details). The dissolved ^15^N_2_ content in the water was measured with a membrane inlet mass spectrometer (PETREL QMS, Bay Instruments), and the percentage enrichment was calculated. At the end of the incubation, the samples (40–1000 mL) were filtered onto a pre-combusted GF/F (450 °C, 4.5 h), thereby ensuring sufficient nitrogen biomass for δ^15^N measurements (>10 µg N and/or >1000 mV amplitude; see the Supporting Information for details). The filters were dried overnight at 60 °C and stored in a desiccator before being packed in tin capsules for mass-spectrometry analyses using an elemental analyzer (Flash 2000 HT, Thermo Scientific, Waltham, MA, USA) coupled to an isotope ratio mass spectrophotometer (Delta V Plus, Thermo Scientific). Three standards (Glutamic Acid USGS 40, Glycine USGS 64, and Caffeine USGS 62) were used for calibration, bracketing the expected δ^15^N range of both enriched and natural abundance samples, and ensuring accuracy. Standards were measured at the beginning, middle, and end of each analytical session (Figure S1). The limit of detection was 0.01 nmol N L^−1^ d^−1^.

#### nifH gene extraction, sequencing, and analysis

Samples (1.5 L) were filtered through a 12 µm (WHA10400012, Whatman, Merck) filter to collect the aggregate-associated microbes, and the filtered water samples were filtered again on a Supor 0.2 µm filter (GPWP04700, Millipore, Merck) to collect the free-living cells. Filters were placed in cryo-tubes with lysis buffer (1 mL; see the Supporting Information for details) and stored at −80 °C until extraction. DNA was extracted by phase separation, as described previously (*25*), and kept in DNAase-free water. *nifH* genes were amplified via nested polymerase chain reaction (PCR; Life ECO, Bioer, Hangzhou, China) with *nifH* 3,4 and *nifH* 1,2 primers, as described in the Supporting Information (*26*). *nifH* samples were sequenced on a <u>sequencing platform (MiSeq, Illumina, San Diego, CA, USA)</u> <u>by RTL Genomics (Lubbock, TX, USA)</u>. *nifH* sequences were analyzed with the Quantitative Insights Into Microbial Ecology 2 pipeline (QIIME2, version 2022.8) (*27*). Amplicon sequence variants (ASVs) were then assigned to *nifH* reference samples (*19*). Before analysis, ASVs representing 5% or less of the total assigned samples at the family level were removed from the final dataset. Samples were grouped based on shared properties (free-living vs. aggregate-associated, summer vs. winter, and river vs. lake). The Shannon index was calculated and compared using the Kruskal–Wallis test with a sampling depth of 1000 ASVs to determine the difference in alpha diversity between groups of samples. For beta diversity, Bray–Curtis distances between samples were generated and compared using analysis of similarities (ANOSIM) to test for significant differences in community structure between groups (*28*).

#### Visualizing diazotrophs by nitrogenase immunolocalization

Following 48 h of incubation, subsamples (50 mL) were filtered by using a gentle pressure (<1 mbar) on a 0.4 µm polycarbonate filter. Subsamples were fixed overnight with chilled ethanol (5 mL). Cell membranes were permeabilized, immunolabeled, and imaged by confocal laser scanning microscopy, as described in previous studies (*29*, *30*).

#### Diazotrophs and bacterial counts by flow cytometry

Water samples (1.7 mL) were fixed with glutaraldehyde for 10 min at room temperature (0.02% final concentration), snap-frozen in liquid nitrogen, and stored at −80 °C until analysis. In the lab, the samples were fast-thawed, mixed with ethylenediaminetetraacetic acid (5 µM, pH 8), and probe sonicated to dislodge the aggregated cells (amplitude 55 mV, 10 s × 3). Cyanobacterial and picoeukaryotic algae were run unstained (300 µL). For total BA measurement, subsamples (100 µL) were stained with SYBR Green for 10 min in the dark. DA was counted using a nitrogenase immunolabeling approach (*31*). Briefly, subsamples (1 mL) were centrifuged (4000 × *g*, 10 min) and washed three times with phosphate-buffered saline–Triton (PBST). Samples were treated with a primary antibody, anti-*nifH (*AS01 021A, Agrisera Antibodies, Vännäs, Sweden) in PBST-bovine serum albumin. After incubation for 1 h at room temperature, samples were washed three times as described above. A secondary antibody (A-11039, Thermo Fisher Scientific) conjugated to a green fluorophore was added. Subsamples were incubated for 45 min, washed three times, and resuspended in phosphate-buffered saline before measurement. The stained, unstained, and immunolabeled samples were run on a flow cytometer (Attune NxT, Thermo Fisher Scientific) at 25 µL min^−1^. Cell-specific N_2_ fixation was calculated by dividing the N_2_ fixation rates by the number of diazotrophic cells.

#### Statistical analyses

The data distribution was determined using the Shapiro– Wilk test. Differences between sites and fractions within a season (summer or winter) were determined using a one-way analysis of variance (ANOVA) and Fisher’s least significant difference (LSD) means comparison (*n* = 12). An additional *t*-test was performed to test for significant difference between summer and winter averages, as well as between the river and lake. A Pearson correlation matrix was used to determine statistical relationships between PP or BP and N_2_ fixation. An additional Pearson test was conducted for free-living bacteria and aggregate-associated diazotrophy (N_2_ fixation rates, abundance, richness, and Shannon Index) to identify relationships with environmental factors (Chl *a*, temperature, C:N, TOC, particle concentration, DIN, TN, TEP, and PO_4_³⁻). A Mantel test was used to assess the relationship between the community composition of free-living and aggregate-associated diazotrophs and the environmental factors listed above. Canonical correspondence analysis was performed to determine the effects of environmental parameters on diazotroph communities of two separate diazotroph models. The statistical significance of the observed patterns was assessed by ANOVA. All the statistical calculations with ASVs were performed with the phyloseq and Vegan R packages (*32*, *33*). All the calculations were performed with a confidence level of 95%.

## Results and Discussion

### Physicochemical variability along the Jordan River and in Lake Kinneret

The Upper Jordan River flows through ~44 km of rural/agricultural land and is the main source of freshwater and nutrients for Lake Kinneret (*34*). The discharge rate of the Jordan River into Lake Kinneret during summer was estimated as ~7 m^3^ s^−1^ (Table 1). This estimate was derived from measurements at the midstream site, as there was no significant loss or gain of water measured in the downstream section of the river. There was a temperature gradient along the river, increasing from the source towards the lake. Turbidity, number of particles (size range of 150–2200 µm), and TEP also increased toward the lake (Table 1). The increasing number of particles was attributed to organic matter, as the flow conditions do not mobilize inorganic particles of >150 µm in size. The increasing number of organic particles originated mostly from the biodegradation of plant litter, micro-macro algae, and other detritus, which increase in abundance during the southeastern Mediterranean summer due to high temperatures and lack of rain. This biotic activity can increase the algal biomass and particle formation by shredder activity (*35*). The concentrations of Chl *a* (phytoplankton proxy) and TOC increased by 10× and 58%, respectively, from the upstream site to the downstream site. In addition, Chl *a* and TOC peaked in the lake water (2.7 µg L^−1^ and 5.1 mg L^−1^, respectively). TN increased gradually from the upstream site to the midstream and downstream sites (2.2 to 2.7 mg L^−1^), although it was lower in the lake water. Correspondingly, the organic carbon C_org_:N_org_ ratio (i.e., TOC:TON) along the river was ~1 and much lower than the value of ~9:4 obtained for the lake water (mol:mol, Table 1). Similarly to TN, the highest DIN concentrations were measured at the midstream and downstream locations, dominated by nitrate (1.6–2.2 mg L^−1^). The orthophosphate concentrations were similar along the river–lake continuum (<0.05 mg L^−1^). Therefore, corresponding N_inorg_:P_inorg_ ratios were well above the Redfield ratio (16:1) along the river but were close to the Redfield ratio in the lake water owing to the relatively low nitrate concentrations (Table 1).

**Table 1.** Physicochemical data for surface water along the Jordan River and in Lake Kinneret.

| Sampling location | Upstream |  | Midstream |  | Downstream |  | Lake |  |
| --- | --- | --- | --- | --- | --- | --- | --- | --- |
| Season<br>Parameter | Summer | Winter | Summer | Winter | Summer | Winter | Summer | Winter |
| Discharge<br>($\text{m}^3 \text{s}^{-1}$) | $1 \pm 0.1$ | $1.3 \pm 0.0$ | $6.9 \pm 0.4$ | $12.9 \pm 4.9$ | Not<br>measure<br>d | Not<br>measure<br>d | - | - |
| Temperature<br>($^{\circ}\text{C}$) | $18.6 \pm 0.6$ | $15.5 \pm 0.7$ | $24.1 \pm 2.7$ | $16.6 \pm 0.6$ | $25.8 \pm 2.9$ | $17.2 \pm 0.1$ | $31.8 \pm 1.1$ | $19.2 \pm 0.2$ |
| Turbidity<br>(NTU) | $3.3 \pm 1.3$ | $5 \pm 4.7$ | $20.1 \pm 3.5$ | $8.9 \pm 1.1$ | $29 \pm 3$ | $9.6 \pm 1.3$ | $2.6 \pm 0.9$ | $1.9 \pm 0.01$ |
| Particles, >150<br>$\mu\text{m}$<br>($\times 10^3 \text{L}^{-1}$) | $3.7 \pm 2.6$ | $2.3 \pm 1.1$ | $35 \pm 25$ | $2.9 \pm 1.5$ | $64 \pm 4.4$ | $3.3 \pm 1.2$ | $25 \pm 9.2$ | $1.4 \pm 1$ |
| Chl a<br>($\mu\text{g L}^{-1}$) | $0.2 \pm 0.1$ | $0.3 \pm 0.1$ | $1.9 \pm 0.5$ | $1.2 \pm 0.4$ | $1.9 \pm 1.6$ | $1 \pm 0.0$ | $2.7 \pm 0.1$ | $3.1 \pm 0.2$ |
| TEP<br>($\mu\text{g GX L}^{-1}$) | $163 \pm 71$ | $75 \pm 20$ | $508 \pm 141$ | $492 \pm 300$ | $624 \pm 277$ | $556 \pm 143$ | $590 \pm 99$ | $583 \pm 164$ |
| TOC<br>($\text{mg L}^{-1}$) | $1.9 \pm 0.3$ | $1.3 \pm 0.4$ | $2.4 \pm 0.1$ | $3.7 \pm 0.3$ | $2.8 \pm 0.1$ | $3 \pm 0.3$ | $5.1 \pm 0.3$ | $5.5 \pm 0.1$ |
| TN<br>($\text{mg L}^{-1}$) | $2.2 \pm 0.2$ | $2.0 \pm 0.3$ | $2.7 \pm 0.4$ | $2.5 \pm 0.2$ | $2.7 \pm 0.2$ | $2.5 \pm 0.3$ | $1.5 \pm 0.1$ | $1.1 \pm 0.3$ |
| $\text{C}_{\text{org}}:\text{N}_{\text{org}}$<br>(mol:mol) | $1 \pm 0.2$ | $1.2 \pm 0.6$ | $1.1 \pm 0.1$ | $1.8 \pm 0.3$ | $1.3 \pm 0.2$ | $1.4 \pm 0.0$ | $9.4 \pm 72$ | $6 \pm 1.3$ |
| $\text{NO}_3^- - \text{N}$<br>($\mu\text{g L}^{-1}$) | $1.2 \pm 0.07$ | $1.2 \pm 0.01$ | $1.9 \pm 0.32$ | $2.1 \pm 0.06$ | $1.8 \pm 0.22$ | $2.1 \pm 0.02$ | $0.04 \pm 0.01$ | $0.19 \pm 0.08$ |
| $\text{NH}_4^+ - \text{N}$<br>($\mu\text{g L}^{-1}$) | $0.03 \pm 0.01$ | $0.02 \pm 0.01$ | $0.03 \pm 0.02$ | $0.03 \pm <0.01$ | $0.03 \pm 0.02$ | $0.02 \pm 0.01$ | $0.02 \pm 0.01$ | $0.11 \pm 0.05$ |
| $\text{PO}_4^{3-} - \text{P}$<br>($\mu\text{g L}^{-1}$) | $0.02 \pm 0.01$ | $0.01 \pm <0.01$ | $0.03 \pm 0.01$ | $0.05 \pm <0.01$ | $0.03 \pm 0.01$ | $0.03 \pm 0.01$ | $0.01 \pm <0.01$ | $0.01 \pm <0.01$ |
| $\text{N}_{\text{inorg}}:\text{P}_{\text{inorg}}$<br>(mol:mol) | $98 \pm 26$ | $165 \pm 7$ | $91 \pm 19$ | $71 \pm 9$ | $84 \pm 5$ | $119 \pm 42$ | $15 \pm 7$ | $50 \pm 27$ |

During winter, average discharge from the Jordan River into Lake Kinneret increased by more than 50% relative to summer (Table 1). The discharge flux was more variable than during summer because of the intense but sporadic rain events (Table 1). The water temperature remained constant along the river (~17 °C) and lower (*P* < 0.01) than in summer (Table 1). Turbidity and particle count were similar along the river (~8 NTU and ~2.4 × 10^3^ L^−1^, respectively), and significantly lower than in the lake (*P* = 0.04 and 0.4, respectively). The numbers of particles were lower because of flow events during winter, which are intense and usually occur every 1 or 2 weeks. These rain events act as reset periods (*35*) that flush out particles and biota from the river toward the lake, thereby preventing biological activity from generating large numbers of particles or the accumulation of particles along the river. Correspondingly, Chl *a* and TEP were lower along the Jordan River than in Lake Kinneret (*P* < 0.01). In contrast to summer, TOC and TN concentrations were similar along the river (3–5 mg L^−1^), with an average C_org_:N_org_ ratio of 1.3 mol/mol. In contrast, TOC concentrations in the lake were higher than in the river, whereas TN concentrations were 50% lower, leading to a C_org_:N_org_ ratio of 5.5 mol/mol (Table 1). Although no trends in DIN and orthophosphate concentrations were detected along the river–lake continuum during winter, N_inorg_:P_inorg_ ratios were much higher than the Redfield ratio, similar to summer.

The ecological stoichiometry along the Jordan River was similar during summer and winter, with N:P values well above the Redfield ratio (*36*) and C:N values well under the ratio (Table 1). These constant ratios were probably due to the ongoing agricultural inputs, comprising various N species, into the river (*37*) and microbial consumption of organic matter. During summer, N:P and C:N ratios in Lake Kinneret were similar to the Redfield ratio, as reported previously (*38*). In Lake Kinneret, these ratios were higher during winter than summer because high external inputs of DIN affect the chemical stoichiometry (P concentration during both seasons was ~0.01 µg L^−1^) of this monomictic environment (*39*, *40*). In contrast to the high variability in C:N:P, the surface water of the Jordan River and Lake Kinneret are rich in iron and molybdenum (*37*, *41*).

Although the physicochemical properties of the surface water along the river– lake continuum were highly variable, the abiotic conditions were generally advantageous for diazotrophs. Specifically, the high availability of trace metals, such as iron and molybdenum, as well as high C:N, which leads to N limitations for heterotrophic bacteria, are reported to encourage diazotrophy (*1*).

### Spatiotemporal changes in diazotroph abundance

The total number of unicellular diazotrophs along the river–lake continuum ranged from ~1 × 10^7^ to ~2 × 10^8^ cells L^−1^ (Figure 2). The number of diazotrophs was based on the number of cells that synthesized nitrogenase, and thus probably fixed N_2_ (*31*). During summer, average diazotrophs along this freshwater continuum accounted for 1%–8% of total BA (Figure 2A). Diazotrophs numbers increased downstream along the river (from 1.5 × 10^7^ to 18 × 10^7^ cells L^−1^), whereas the lowest counts were measured at the lake (0.97 × 10^7^ cells L^−1^). Heterotrophic diazotrophs comprised 29% of the total diazotrophs (phototrophs + heterotrophs) at the upstream site, 36% at the downstream site, and 55% at the lake (Figure 2B). Within the heterotrophic fraction, the highest numbers of aggregate-associated diazotrophs were recorded at the midstream and downstream sites (from 9.1 × 10^6^ to 11.2 × 10^6^ cells L^−1^), whereas the free-living fraction was dominant at the upstream and lake sites (Figure 2C).

**Figure 2.**
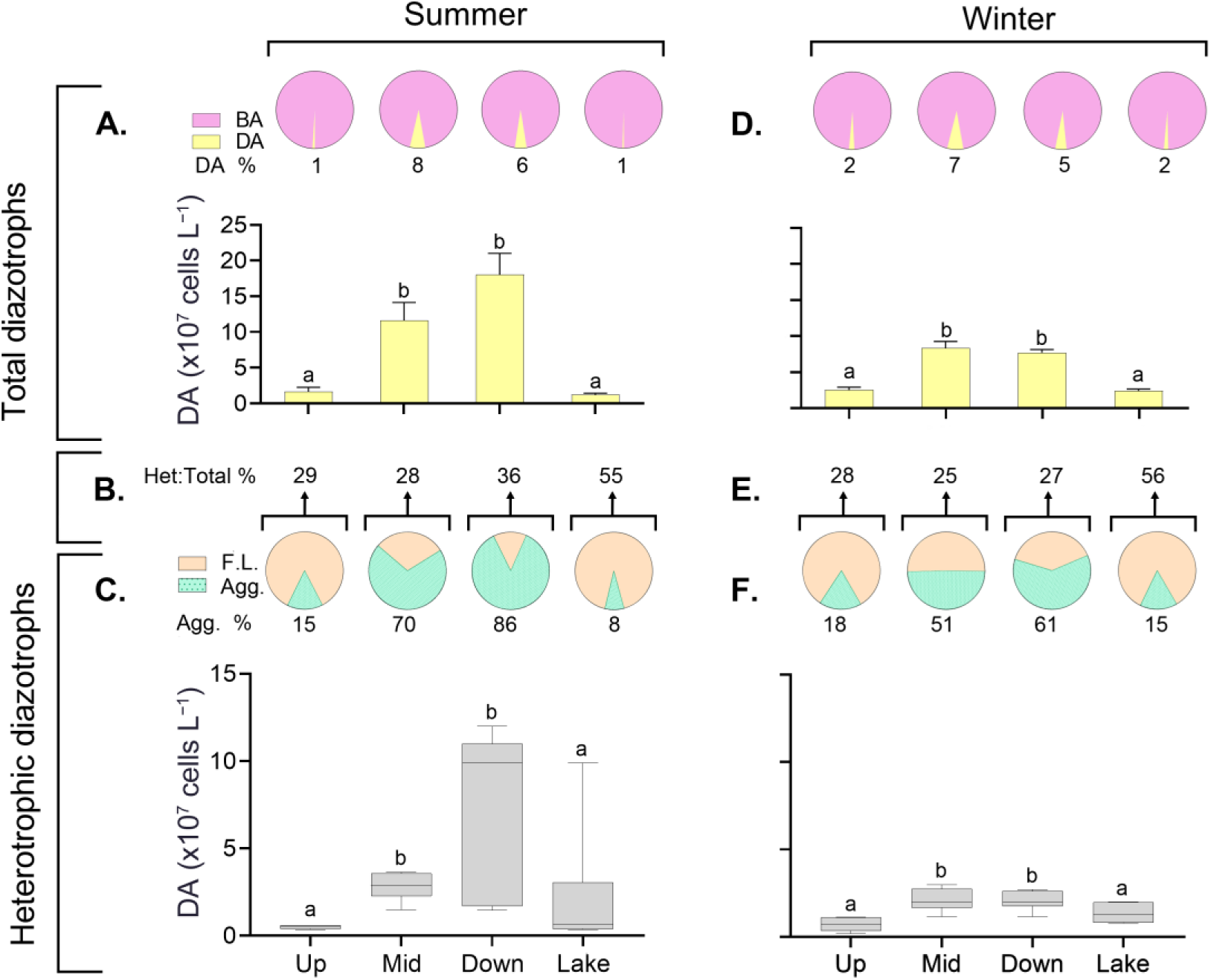
Total DA along the Jordan River and Lake Kinneret in (A) summer and (D) winter. The upper pie bar charts (pink and yellow) represent DA (pink) as a percentage of total BA (yellow). Grey box plots represent the total heterotrophic DA in (C) summer and (F) winter. The lower pie bar charts (orange and green) represent DA associated with aggregates as a percentage of total heterotrophic diazotrophs (free-living and aggregate-associated shown in orange and green, respectively). The percentages represent the fraction of total heterotrophic diazotrophs among total diazotrophs in (B) summer and (E) winter. Lowercase letters indicate significant difference between samples within a season (ANOVA, Fisher’s LSD posthoc test). F.L.: free-living; Agg.: aggregate-associated; Het:Total %: fraction of total heterotrophic diazotrophs among total diazotrophs.

During winter, diazotroph abundance was 2.5 × 10^7^ to 8.2 × 10^7^ cell L^−1^, accounting for 2%–7% of the total bacterial numbers (Figure 2D). The highest numbers of diazotrophs were found at the midstream and downstream sites (7.7 × 10^7^ to 8.3 × 10^7^ cell L^−1^). Nevertheless, the diazotroph abundance was lower than in summer (31% and 2 times, respectively, Figure 2D). The heterotrophic diazotrophs made up 25%–28% of the total N_2_ fixers along the river sites and 56% of those sampled in the lake (Figure 2E). Similar to summer, aggregate-associated diazotrophs increased along the river in winter (2.1 × 10^6^ to 9.9 × 10^6^ cell L^−1^), although the free-living fraction dominated the surface water (8.7 × 10^6^ to 1 × 10^7^ cell L^−1^) at the upstream and lake sites (Figure 2F).

Little is known about the number of diazotrophs across the freshwater ecospace (*10*). Overall, the diazotrophs fraction of the total BA was constrained to <8% along this freshwater ecosystem because of competition for phosphate (*42*) as well as high concentrations of DIN, dissolved oxygen, and increased ratios of N:P that impair N_2_ fixation (*1*). The former is suggested since other abiotic factors that often limit diazotrophy, such as low water temperature, limiting concentrations of available carbon, and low concentrations of iron or molybdenum concentrations, are high in this freshwater ecosystem (*37*, *41*).

Aggregates encourage heterotrophic diazotrophy under conditions unfavorable for diazotrophy (*43*). The aggregates act as focal points for colonization (*44*) which encourages heterotrophic diazotrophy by providing a labile carbon source while simultaneously inducing DIN limitation. During summer, the concentrations of large particles, including TEP, along the Jordan River were highest at the midstream and downstream sites, leading to a clear trend in heterotrophic diazotrophs associated with aggregates. These aggregates may protect diazotrophs from the drastic changes in abiotic conditions within this fluvial environment. In contrast, during winter, low numbers of particles along the river reduced the likelihood of bacterial colonization, decreasing the counts of diazotrophs associated with aggregates. In Lake Kinneret, the high C_org_ concentration, which is mostly dissolved (*45*), and the relatively high C:N ratios could support the metabolic requirements of free-living heterotrophic diazotrophs (*18*).

### Contribution of heterotrophic diazotrophs to total N_2_ fixation

Total N_2_ fixation rates (i.e., by phototropic and heterotrophic diazotrophs) ranged from 0.1 to 1.9 nmol N L^−1^ d^−1^, increasing downstream along the river and peaking at the lake (Figure 3A, E). The contribution of heterotrophic diazotrophs to total N_2_ fixation rates was highest at the upstream site (100%) and decreased to 19% in the lake (Figure 3B). N_2_ fixation rates for heterotrophic diazotrophs were highest at the midstream and downstream sites (~0.27 nmol N L^−1^ d^−1^), attributed mostly to diazotrophs associated with aggregates (Figure 3C). Overall, the N_2_ fixation rates per cell were higher for diazotrophs associated with aggregates (61.7 amol N cell^−1^ d^−1^) than for the free-living fraction (16.5 amol N cell^−1^ d^−1^). These cell-specific rates decreased along the river and were highest in the lake (Figure 3D).

**Figure 3.**
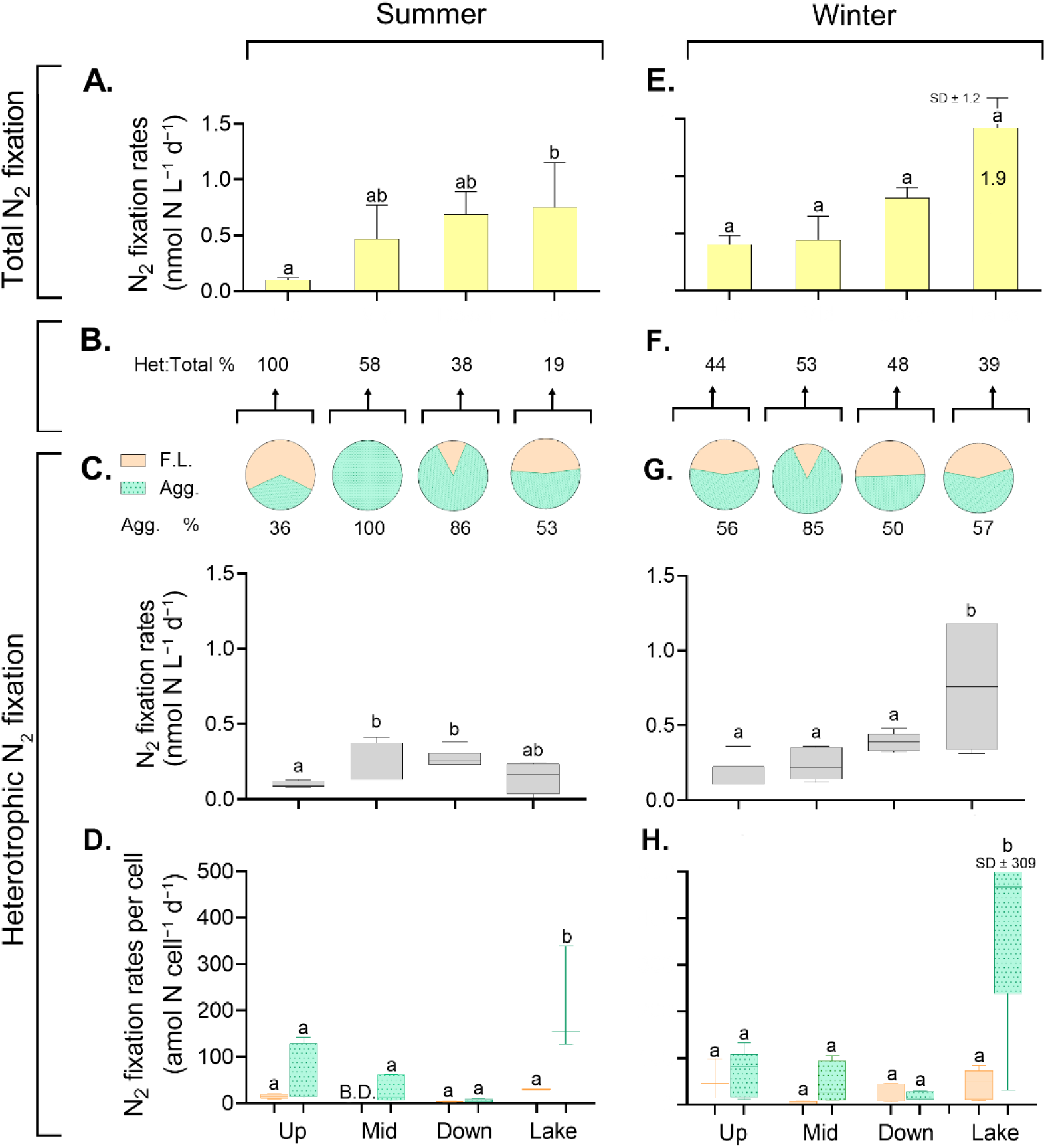
(A–D) Summer and (E–H) winter values of total N_2_ fixation along the Jordan River and Lake Kinneret (A, E), and the percentage contribution of heterotrophic diazotrophs to total N_2_ fixation (B, F). Pie charts show the fraction of heterotrophic N_2_ fixation (100%, gray bar plot) relative to free-living (orange) and aggregate-associated (green) fractions (C, G). Free-living or aggregate-associated heterotrophic N_2_ fixation per cell (D, H) was calculated by dividing the corresponding N_2_ fixation rate by DA. Lowercase letters indicate significant difference between samples within a season (ANOVA and Fisher’s LSD post hoc test). B.D. indicates measurements that were below the detection limit. F.L.: free-living; Agg.: aggregate-associated; Het:Total %: fraction of total heterotrophic diazotrophs among total diazotrophs.

During winter, heterotrophic diazotrophs contributed 39%–53% of total N_2_ fixation with no spatial trend (Figure 3F). Compared with summer, average heterotrophic N_2_ fixation rates were higher by up to 5 times in the river and ~1.5 times in the lake, ascribed equally to both free-living and aggregate-associated diazotrophs (apart from the midstream site) (Figure 3G). Exceptionally high N_2_ fixation rates per cell were ascribed to heterotrophic diazotrophs associated with aggregates in the lake (496 ± 309 amol N cell^−1^ d^−1^), whereas the overall trend was similar to that found in summer (Figure 3H).

Overall, N_2_ fixation rates for heterotrophic diazotrophs have rarely been quantified in freshwater environments and have only been measured sporadically within corresponding sediments (*17*). The contribution of heterotrophic diazotrophs to total N_2_ fixation was high along this fluvial system and moderate in the lake (Figure 3B). Heterotrophic diazotrophs, in contrast to phototrophs, tolerate variable conditions and thus can fix N_2_ at high rates even as the water temperature changes (*46*, *47*). This ability to acclimate swiftly could provide heterotrophic diazotrophs with the capacity to fix N_2_ at high rates in highly dynamic environments such as the Jordan River.

Our results show that heterotrophic diazotrophs along this freshwater continuum were fixing N_2_ at high rates despite conditions that should suppress this process, including high oxygen saturation, DIN concentrations, and N:P ratio (*43*, *48*). N_2_ fixation rates per cell for aggregate-associated diazotrophs were generally higher than those for free-living cells. The N_2_ fixation efficiency for aggregate-associated heterotrophic diazotrophs may be higher than that of free-living cells because the organic complexes in the aggregates provide physicochemical advantages that encourage diazotrophy (*43*). Large aggregates (>0.5 mm) can create oxygen-minimum microzones that may protect the nitrogenase from ambient oxygen (*48*). In addition, aggregates, compared with their surroundings, provide microenvironments with higher availability of labile carbon, which is required to sustain the high metabolic needs of N_2_ fixation as the various prokaryotes that colonize the aggregate degrade the organic matrix (*4*, *49*). Furthermore, aggregate-associated microbes typically remineralize nitrogen species at higher rates than carbon species, thereby increasing the C:N ratio of the aggregate (*50*), creating a nitrogen-limited microenvironment and further encouraging diazotrophy.

### Diazotroph lifestyle: community composition of free-living or aggregate-associated cells

Amplicon sequencing analysis indicated high spatiotemporal variability in the community, diversity, and richness along the river–lake continuum between free-living and aggregate-associated diazotrophs (Figure 4A, B). At the upstream site, free-living non-cyanobacterial diazotrophs (NCDs) were dominated (30%) by Nitrosomonadales (Betaproteobacteria) (Figure 4A). Free-living NCDs at the mid-stream site were mostly assigned to Desulfuromonadales (17%) and at the downstream site by Methylococcales (30%), whereas Vibrionales dominated the NCDs in the lake (39%). Some major diazotroph families, such as Nitrosomonadales and Desulfuromonadales, were found in the oxygenated river despite being obligate anaerobes (Figure 4A). According to DNA-*nifH* sequencing, their presence was attributed to anaerobic diazotrophs associated with biofilms attached to the streambed that were resuspended under the rapid flow. However, the N_2_ fixation rates of these oxygen-sensitive diazotrophs were probably decreased because they were free-living cells in an oxygenated environment and oxygen irreversibly damages nitrogenases (*51*). This possible decrease is consistent with N_2_ fixation rates, indicating that the majority of N_2_ fixation in the midstream and downstream sites could be attributed to aggregates (Figure 3C, D). At Lake Kinneret, *Cylindrospermopsis* accounted for about half of the N_2_-fixing bacteria found during summer and was dominant in winter (Figure 4A). *Cylindrospermopsis* is a filamentous cyanobacterial diazotroph that forms strings of cells of 10–200 µm in length with a nominal diameter of <5 µm (*52*). This diazotroph is ubiquitous and can fix N_2_ under changing physicochemical conditions (*53*). *Cylindrospermopsis* can form intensive blooms in a range of lakes (*52*, *54*) and has exceptional physiological adaptability, allowing it to invade a wide range of new freshwater environments (*55*).

**Figure 4.**
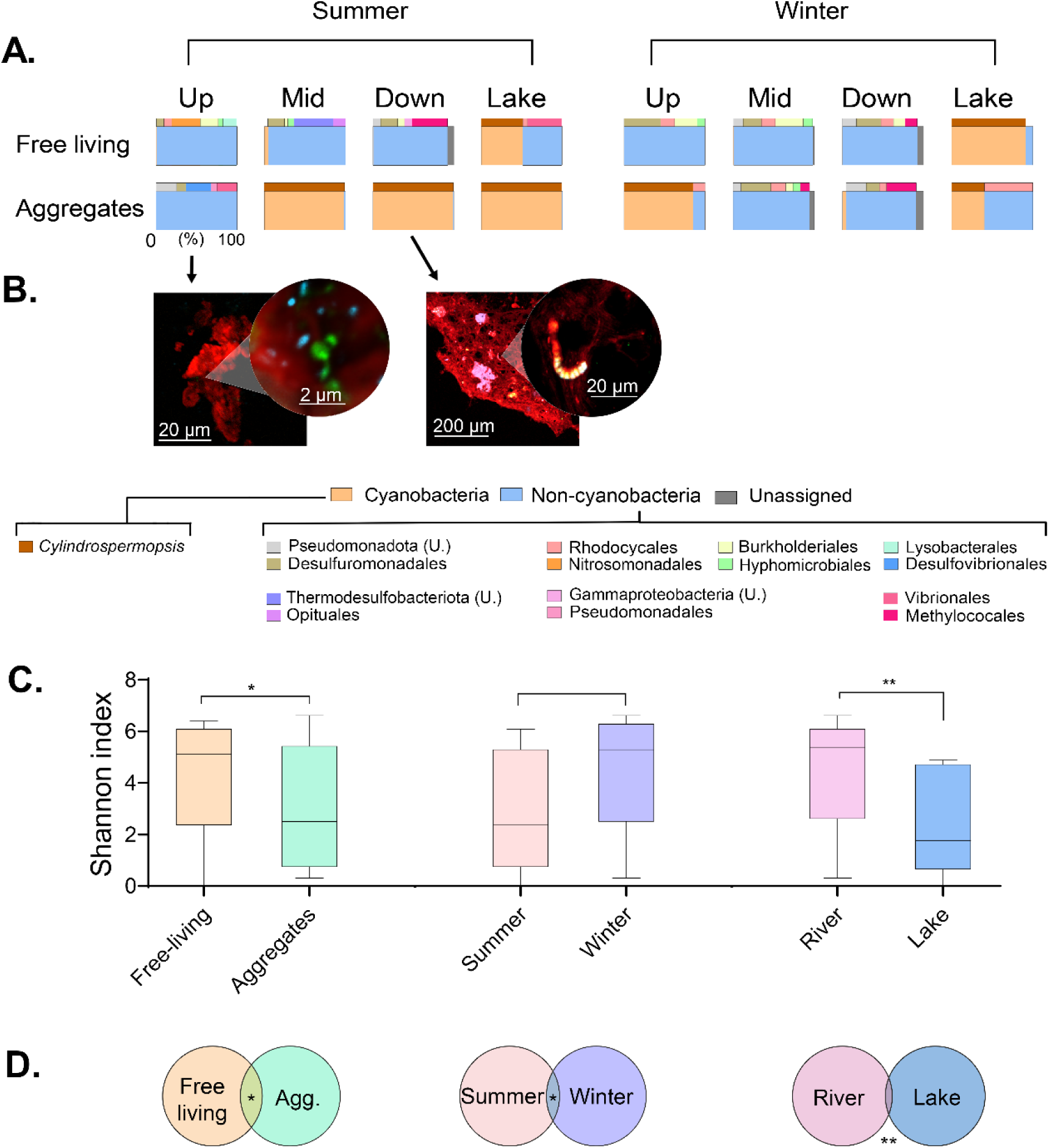
(A) Fractions of cyanobacterial diazotrophs (genus) and NCDs (orders) as well as unassigned ASV sequences along the river–lake continuum retrieved during summer and winter. Bar charts above the fractions represent the families (>5%) of total NCDs (>10% from total ASV sequences). U. represents unassigned orders of phyla. (B) Representative confocal laser scanning microscope images of immunolabeled diazotrophs that were collected from the Jordan River at the upstream and downstream sites. Micrographs were taken in four layers, showing diazotrophs by immunolabeling of the nitrogenase (green), cyanobacteria by phycoerythrin autofluorescence (light orange), bacteria by DAPI staining (light blue), and the polysaccharide matrix of the aggregates by Concanavalin A staining (red). Insets show magnified regions of interest (note the different scale bar). (C) Alpha diversity was determined for all ASV sequences based on shared properties and compared for free-living vs aggregate-associated, summer vs winter, and river vs lake diazotrophs. (D) Beta-diversity was calculated for all the ASVs described in B and compared by applying an ANOSIM test. Overlapping areas indicate the number of ASV sequences that were found in both samples. Asterisks indicate the significance: * *P* < 0.05; ** *P* < 0.01.

Unlike the free-living fraction, the diazotroph community associated with aggregates was dominated by *Cylindrospermopsis* (98%–99%) during summer (Figure 4B), except from the upstream site. *Cylindrospermopsis* often secretes organic matter as polysaccharides (*56*), and thus can act as an active hub for aggregation and bacterial colonization. At the upstream site, aggregate-associated NCDs (Figure 4B) were assigned to Desulfovibrionales (27.9%), Vibrionales (22.2%), and Pseudomonadota spp. (23.1%, unassigned genera). Despite the prevalence of *Cylindrospermopsis* in the analyses, the richness of diazotroph species in the aggregates was similar to that of the free-living fraction. However, a significant dissimilarity between size fractions was attributed to diazotrophic species, such as Desulfovibrionales or Pseudomonadales, which were only associated with aggregates (P = 0.04, Figure 4C, D). Aggregates at the upstream site were solely colonized by NCD. During summer, diazotrophs were less diverse (2.7, Shannon index) than during winter (4.5), with a 40% lower richness value (P < 0.05). The significant dissimilarity between seasons (Figure 4C, D) was attributed to physicochemical changes (Figure 5) that lead to shifts in the proportions of various diazotrophs, such as Vibrionales, which were only observed during summer. Although free-living NCDs continued to dominate this fluvial system (93%–98%), the lake samples were dominated (91%) by *Cylindrospermopsis* (Figure 4A). Free-living NCDs in the river were dominated by Alphaproteobacteria and Betaproteobacteria (Figure 4A), including Hyphomicrobiales (12%) and Burkholderiales (20%) which were previously found associated with various plants (57). We hypothesized that high flow rates during the winter could resuspend these diazotrophic phylotypes into the stream with plant litter washed from the riverbanks. The high abundance of these NCDs could also explain the higher numbers of free-living diazotrophs and N_2_ fixation rates measured along the river during winter (Figure 2E, F).

**Figure 5.**
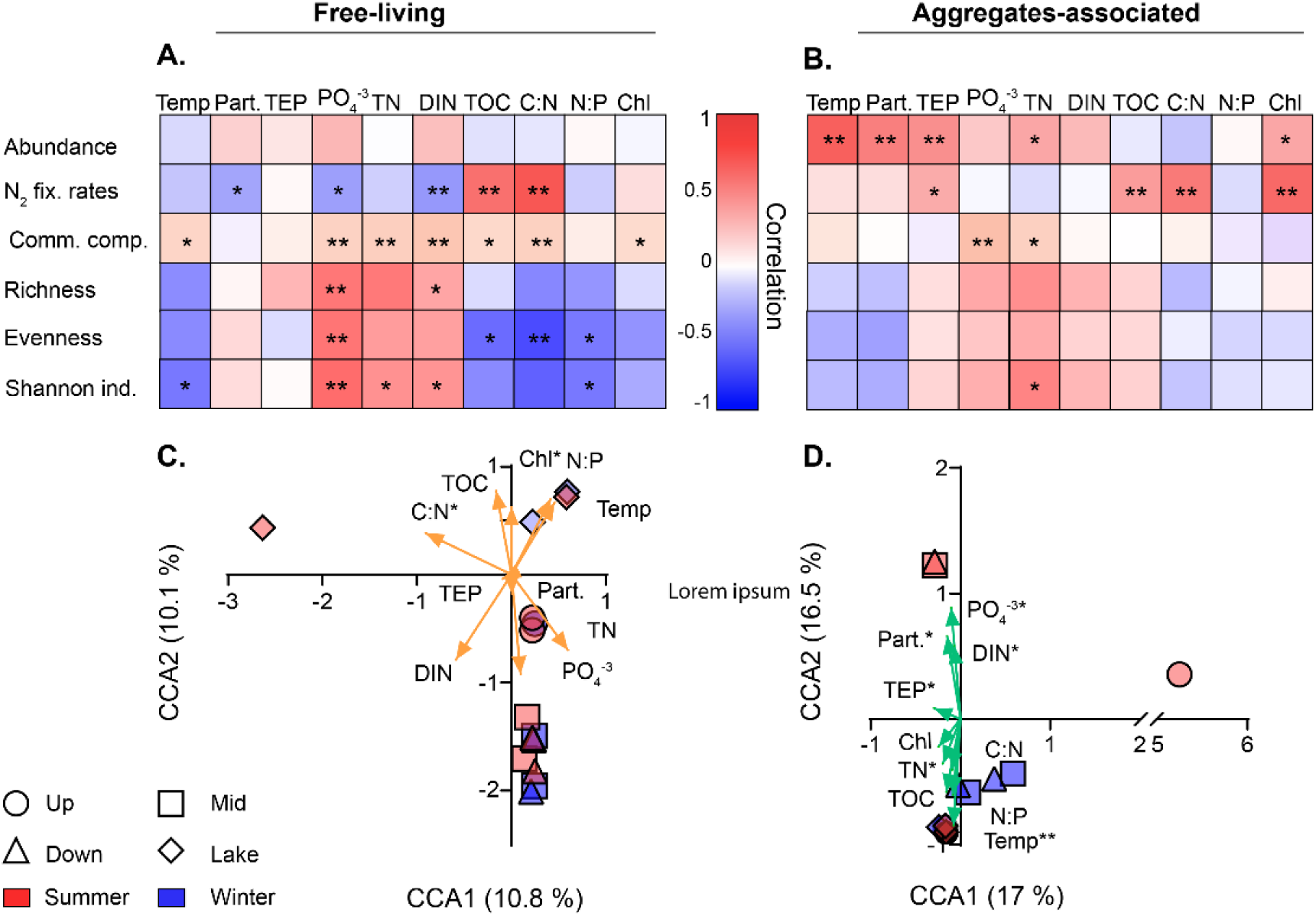
Relationship between environmental drivers and diazotrophy as free-living or aggregate-associated bacteria. N_2_ fixation rates, DA, richness, evenness, and Shannon index were analyzed using a Pearson correlation, and the community composition was compared using the Mentel test for free-living (A) and aggregate-associated (B) bacteria. Beta-diversity was determined by a canonical correspondence analysis model for free-living (C) and aggregate-associated (D) bacteria. Asterisks indicate significance levels: * *P* < 0.05; ** *P* < 0.01.

In contrast to the free-living fraction, aggregate-associated diazotrophs were dominated by *Cylindrospermopsis* at the upstream site (91%), and NCD were also identified on aggregates from the midstream site to the lake site. Three primary diazotrophic groups dominated at each sampling location: Desulfuromonadales (21%) at the midstream site; Methylococcales (42%) at the downstream site; and Rhodocyclales (24%) in the lake. Based on these results, the Jordan River contains a richer diazotroph community than the lake, harboring site-specific diazotrophs, mainly from *Gammaproteobacteria* and *Thermodesulfobacteriota*, including *Vibrionales* and *Opitutales*. The heterogenic nature of the diazotrophic communities found in the Jordan River results from the immense physicochemical variability found in lotic environments compared with lentic environments.

### Effect of environmental factors on diazotrophy along the Jordan River-to-Lake Kinneret continuum

No significant correlation was detected between the abundance of free-living heterotrophic diazotrophs and the environmental factors tested herein (Figure 5A). These findings are consistent with the relatively low numbers of free-living diazotrophs measured along this river–lake continuum (Figure 2C). In contrast, the abundance of diazotrophs associated with aggregates was positively and significantly correlated with increasing water temperature and higher concentrations of large particles (>150 µm), TEP, TN, and Chl *a* (Figure 5B). We suggest that these abiotic conditions encourage microbial colonization of aggregates and facilitate their growth, including diazotrophs. Higher numbers of large particles, such as TEP (polysaccharide-rich and sticky molecules), probably resulted from phytoplankton production, measured by Chl *a* (*58*). These organic particles increase the probability of bacterial colonization (*49*, *50*) and provide a biochemical advantage for diazotrophs in aquatic environments with high TN (*48*).

N_2_ fixation rates for free-living heterotrophic diazotrophs were negatively correlated with increasing particle number and high concentrations of orthophosphate and DIN (Figure 5A). It is unclear how and why particles impair N_2_ fixation rates for free-living diazotrophs, although this correlation could be indirectly linked to competition with aggregate-associated diazotrophs for trace metals and other nutrients. Although increased orthophosphate concentrations were expected to promote diazotrophy (*59*), the negative correlation reported here may have been related to the high DIN concentrations and high N:P ratio (Table 1). Correspondingly, high DIN concentrations along this freshwater continuum may impair the nitrogenase activity and lead to these low N_2_ fixation rates, as reported for other ecosystems (*60*, *61*). In contrast, TOC and high C:N ratios were positively correlated with N_2_ fixation by free-living heterotrophic diazotrophs (Figure 5B). Increased TOC concentrations and C:N ratio can increase N_2_ fixation rates by meeting the metabolic requirements of nitrogenase while inducing N limitation (*62–64*).

N_2_ fixation by aggregate-associated heterotrophic diazotrophs was positively correlated with increased TEP concentration, C:N ratio, and Chl *a* concentration (Figure 5B). These factors can support the colonization of bacteria on aggregates (*65*, *66*) and provide advantageous conditions for diazotrophs to fix N_2_ (*43*, *67*). Unlike free-living heterotrophic diazotrophs, DIN and orthophosphate did not affect N_2_ fixation rates for aggregate-associated diazotrophs (Figure 5B). Despite changes in abiotic conditions in the water column, aggregates, especially the larger ones (>500 µm), may provide beneficial microzones for heterotrophic diazotrophs (*48*) and partly protect them from the surrounding aquatic environment.

The community composition of free-living diazotrophs was positively correlated with most environmental variables (Figure 5A). Conversely, the diazotroph richness increased only with orthophosphate and DIN concentrations (Figure 5A). The corresponding evenness also increased with orthophosphate, yet was negatively correlated with TOC, C:N, and N:P ratios (Figure 5A). Shannon index values were positively correlated with orthophosphate and nitrogen species, and negatively correlated with increasing water temperature and N:P ratio (Figure 5A). These results indicate that water eutrophication (mainly N and P) could increase the diversity of free-living diazotrophs, as previously reported for other freshwater bodies (*68*). In contrast to the free-living fraction, the diversity of aggregate-associated diazotrophs increased only with increasing concentrations of orthophosphate and TN (Figure 5B). We suggest that increased eutrophication not only increased the diversity of free-living diazotrophs, but also supported particle colonization (*69*), thereby increasing the diversity of aggregate-associated diazotrophs indirectly.

The environmental factors that affect the spatiotemporal distribution of diazotroph beta-diversity were analyzed further by canonical correspondence analysis (Figure 5C, D). The diversity of free-living diazotrophs was positively correlated with Chl *a*, increased N:P ratio, and increased water temperatures in the lake during summer and winter (Figure 5C). *Cylindrospermopsis* was clustered with these lake samples in both seasons (Figure S2). This invasive cyanobacterial diazotroph fixes N_2_ under a wide range of water temperatures and N:P ratios (*54*). Nevertheless, an increased C:N ratio was strongly linked with *Vibrio* sp. recovered during summer from the lake. Heterotrophic diazotrophs, such as *Vibrio natriegens* and *Vibrio diazotrophicus* (*70*), are favored by an increased C:N ratio that supports metabolic activity and induces nitrogen limitation (*4*, *18*, *61*). In contrast, the trend in river diazotroph diversity was affected mainly by nutrient loads rather than water temperature, as indicated by the tight clustering of the pristine upstream site (Figure 5C).

In contrast, the beta-diversity of diazotrophs associated with aggregates found at the midstream and downstream sites was mostly linked to increased eutrophication (Figure 5D), which is persistent in the river due to agricultural inputs from the Hula Valley (*37*) especially during summer when water flows are low (Table 1). The aggregates at these sites comprised mainly *Cylindrospermopsis* (Figure 4A, B). These cyanobacterial diazotrophs can attach to other organic particles and aggregate (*71*) or form hubs for colonization because they secrete sticky polysaccharides (*72*). Conversely, the diversity of diazotrophs associated with aggregates clustered at the midstream, downstream, and lake sites during winter (Figure 5D). These diazotroph clusters were mostly assigned to NCDs (Rhodocyclales [20%–100%], and, Desulfuromonadales [19%–39%], and were strongly affected by water temperature. Clustering of aggregate-associated NCDs in the river samples during winter, or during both seasons at the upstream site, could also be linked indirectly to the resuspension of aggregates and diazotrophs from sediment (*5*). Therefore, NCDs clustering on aggregates may be found year-round in locations where flow is generally high and particulate organic matter, such as plant litter, is resuspended.

### Contribution of N_2_ fixation to freshwater PP and secondary production

During summer, PP and total heterotrophic BP (sum of free-living and aggregate-associated bacteria; Figure S3) rates increased gradually from the upstream site to the lake (Figure 6A, B). This increase in PP and BP along the fluvial ecosystem was probably caused by the positive gradient in water temperature and increased concentrations of nutrients and labile organic carbon from plant litter transported in the river. In contrast, the reduction in PP and BP along the river during winter was probably caused by lower water temperatures, similar to a previous study that reported BP and PP increased with water temperature (*73*). PP was significantly linked to total N_2_ fixation rates (Figure 6E). In contrast, no correlation was found between heterotrophic N_2_ fixation rates and BP (Figure 6F).

**Figure 6.**
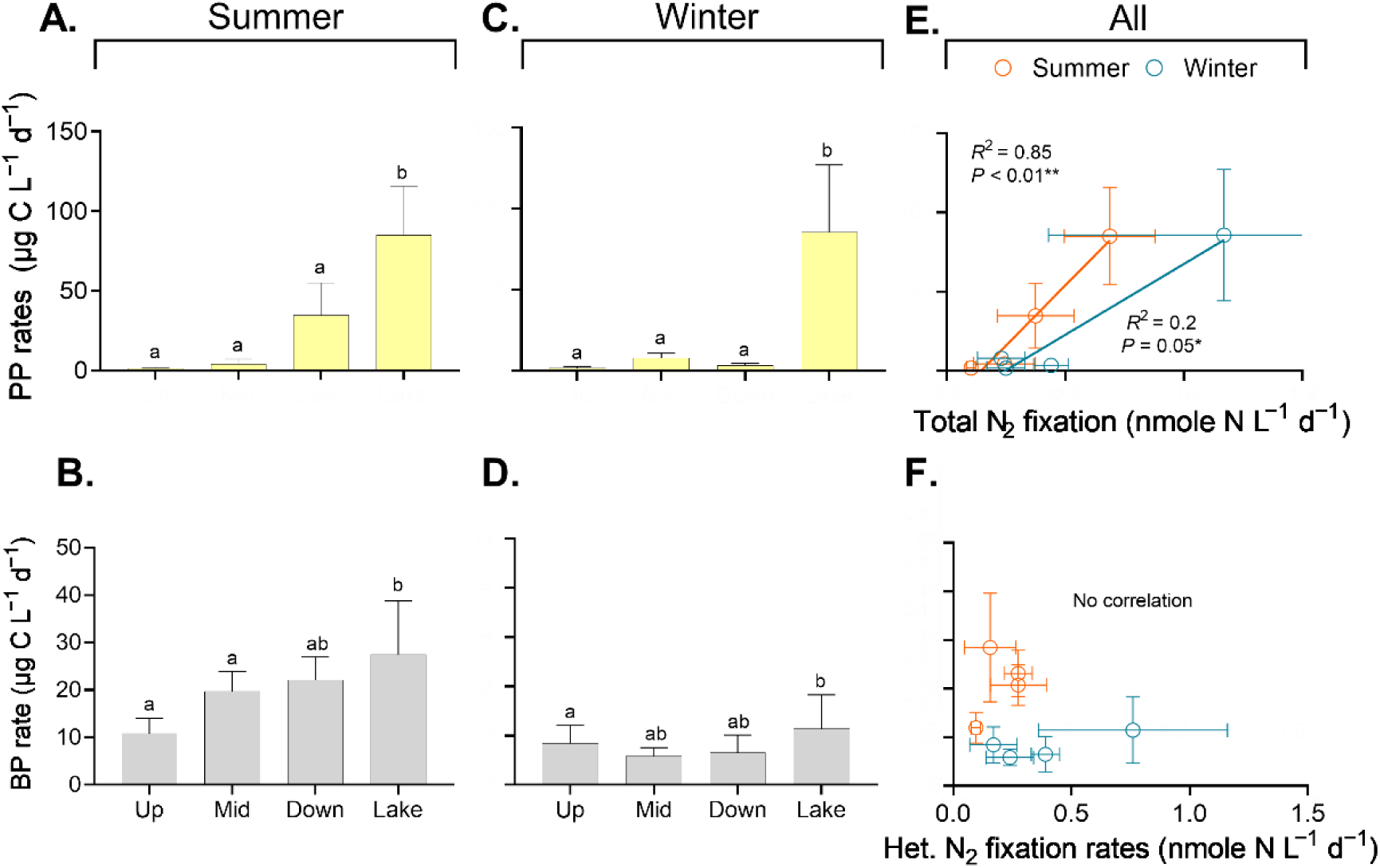
Spatial distribution of PP (A and C) and BP (B and D) along the Jordan River and Lake Kinneret during summer (A and E) and winter (C and D), and a correlations matrix between PP or BP and N_2_ fixation (E and F). Asterisks indicate significance levels: * *P* < 0.05, ** *P* < 0.01.

During winter, no trend was found in the PP or BP rates along the river–lake continuum (Figure 6D, E), probably because of low water temperatures and strong mixing (*74*). A weak, significant correlation was found between PP and total N_2_ fixation, whereas no relationship was observed for BP and heterotrophic N_2_ fixation rates (Figure 6E, F). These results indicate that total N_2_ fixation strongly supported PP in this freshwater environment, as reported previously for marine and terrestrial ecosystems (*75–77*). Moreover, this correlation suggests that nitrogen fixers are important agents for primary producers, not only in oligotrophic environments but also in eutrophic freshwater ecosystems.

## Summary

Little is known about heterotrophic diazotrophy, specifically the abundance, N_2_ fixation rates, and diversity of N_2_ fixers across the freshwater ecospace. Our findings highlight the immense spatiotemporal variability in heterotrophic diazotrophy along the Jordan River-to-Lake Kinneret continuum. Despite this variability, the number of heterotrophic diazotrophs rarely exceeded 10% of the total bacterial community, whereas the fraction of heterotrophic to phototrophic diazotrophs was highly dynamic and ranged from 14.7% to 76.7%. Moreover, the lifestyle of diazotrophs as free-living or aggregate-associated cells affected their abundance, diversity, and contribution to N_2_ fixation. The abundance of aggregate-associated N_2_ fixers may account for up to 86% of all heterotrophic diazotrophs and often contributed more than 50% of the total N_2_ fixation. The diversity of free-living diazotrophs was strongly affected by various environmental drivers, whereas that of aggregate-associated diazotrophs was less susceptible to changing environmental conditions. Finally, the importance of biogeochemical processes associated with aggregates suggests that their inclusion in catchment-scale modeling of nutrients is essential. Based on our results and the contribution of diazotrophs to PP along this river–lake continuum, freshwater diazotrophy and diazotroph lifestyles should be explored further in other freshwater ecosystems.

## Acknowledgments

We thank Dr. Joseane Marques for her assistance with the statistical analysis. This study was supported by the Israeli Science Foundation (grant number 944\21 to E.B-Z). We also thank Adva Speter for assisting with the sampling campaigns.

## Author Contributions

EG, ER, and EB-Z conceived and designed the sampling campaigns and experiments. Material preparation, and experiments were performed by EG. ER and EB-Z, SA. contributed reagents, materials, and analysis tools. GS-V analyzed nutrients samples. HS measured the N_2_ fixation rates together with EG. MK assisted in bioinformatics analyses. The manuscript was written by EG, and EB-Z, whereas all authors commented and contributed to previous versions of the manuscript. All authors read and approved the final manuscript.

