## Supplementary material for "Heterotrophic diazotrophy along a river–lake continuum: lifestyle and contribution to N_2_ fixation"

### Materials and Methods

*Diazotrophs and bacterial counts via flow cytometry.* Subsamples (1.7 ml) were fixed with glutaraldehyde (final concentration 0.02% v:v, Sigma-Aldrich G7651) and incubated for 10 min in the dark. Following 10 min, samples were snap-frozen in liquid nitrogen and stored at -80 °C before measurements. subsamples were then thawed at room temperature, and ethylenediaminetetraacetic acid (EDTA, Sigma Aldrich, 03690, final concentration 5 µM) was added before sonicating subsamples using a probe sonicator (~10 sec, x3 times) for breaking aggregates' matrix and releasing cells. 1000 µl of subsamples were taken for DA measurements while 700 µl were taken for BA.

*Counting diazotrophs by nitrogenase enzyme immunolabeling.* Subsamples were centrifuged (x4000 g, 10 min) and the supernatant was cautiously discarded to maintain bacterial pellet and washed with PBST. The procedure was repeated three times to ensure membrane permeabilization. Fresh anti-nifH (3 µg $ml^{-1}$, final concentration) was prepared with sterile BSA and PBST to increase the specific link efficiency between the antibody and nitrogenase enzyme. Samples were incubated for one hour under room temperature with slow rotation (Benchmark Scientific Roto-Therm Plus, H2024). Subsamples were washed again as described previously. A secondary antibody conjugated to a green fluorophore (Alexa Fluor™ 488) was added to subsamples in the same concentration as the first antibody (Thermo Fisher Scientific A-11039). Subsamples were warped to avoid exposure to the light and incubated for 45 in the same conditions. Residues of secondary antibody were removed by the same washing procedure, and 1 ml of PBS was added to subsamples. A negative control (without first antibody) to verify unspecific adsorption by secondary antibody. Several subsamples were stained (details below) to count the total bacteria and estimate cell loss during the washing procedure (Geisler et al., 2023).

*Quantifying total bacteria.* subsamples were also stained with SYBR Green I (S7563, Invitrogen, final concentration, 1 nM) for 15 min in the dark to count all the bacteria. Separate subsamples were left unstained to determine the phototrophic bacterial concentrations by the autofluorescence of phycoerythrin and chlorophyll *a* pigments.

*Flow cytometry analysis*. Non-stained, stained and nitrogenase-immunolabeled samples were measured with Attune-Next Acoustic Flow Cytometry equipped with a 450 nm laser (blue) detector of 520±30 nm for dye and 574±26 for non-dyed cells. For every five samples, the system was washed with DDW (sterile), and for every 12 samples, beads (1 µm, F8815, Invitrogen, Ex: 350 nm Em: 440 nm) were added to calibrate the size of cells (final concentration, $1.8 \times 10^4$ beads $ml^{-1}$). The flow rate was set for 100 $\mu l^{-1,}$ and the stop condition was $20 \times 10^3$ events per sample.

*Preparation of $^{15}N$ rich water* – One liter of autoclaved phosphate-buffered saline (PBS) was prepared by dissolving one tablet in 200 mL of double-distilled water (P4417, Merck). The solution was degassed overnight using a degassing membrane (G543, MiniModule) connected to a vacuum pump at room temperature. The bottle was then disconnected from the degassing system without exposure to air, and 15 mL of $^{15}N_2$ gas (99 %, Cambridge Isotopes) was added. The bottle was shaken vigorously for one hour and stored at 4°C for a week before the water was added to the microcosm bottles. (1)

*Quality tests of EA-IRMS analyses*. A few measures were taken throughout the measurements to ensure the accuracy and stability of the isotopic analyses. Quartz crucibles were used and replaced every 15-20 samples to allow the removal of the ash and facilitate clean operation of the reactor over time. Accuracy was ensured by a calibration of the measured isotopic values with a set of secondary standards (Figure S2. A), bracketing the range of isotopic values of both natural abundance and enriched samples (Caffeine USGS62, $\delta^{15}N_{AIR\text{-}N2}$=+20.17‰; Glycine USGS64, $\delta^{15}N_{AIR\text{-}N2}$=+1.76 ‰; and L-Glutamic acid USGS40, -4.52 ± 0.06 ‰). Linearity test was performed at the beginning of the work to determine the range of nitrogen levels in the samples, which maintain a constant stable isotopic value using USGS 64 (Figure S2. B). According to the results, a range of 10 to 52 µg N in samples was determined. These values are in accordance with previous publications (2, 3). Standards were measured at the beginning and the end of each set. Also, a representative standard (Glycine USGS64) was measured repeatedly every 4-5 samples to verify the accuracy and correct for a potential drift when needed. No significant drift nor deviation was observed during the measurements (Figure S2 .C, standard deviation = ±0.3‰). Finally, a quantification of N amount in samples' biomass was achieved by a correlation of peak amplitude to known values in the Acetanilide. A linear trend line was captured over a wide range of N masses, enabling reliable measurements of 1.8 to 52.4 µg N in samples (Figure S2. C).

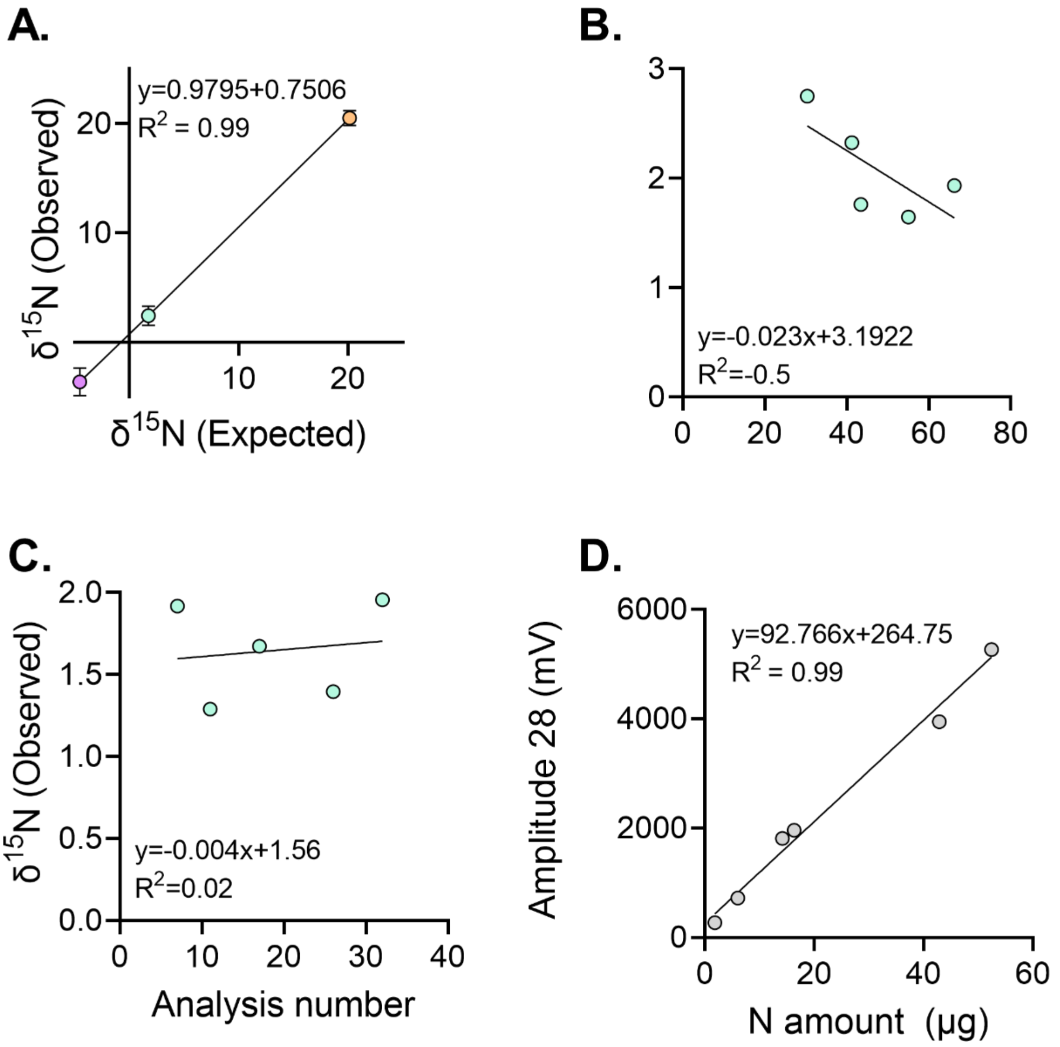


**Figure S1.** EA-IRMS quality control tests. A calibration curve between expected and measured δ$^{15}$N for a range of known secondary standards, bracketing the samples' values (n=32) (A). Linearity test of δ$^{15}$N in Glycine over a range of N amount (µg) in samples (standard deviation for the range of 1.8 to 52.4 µg N was 0.1‰) (B). A drift test of Glycine throughout the analysis showing no significant change (slope = -0.004, and standard deviation ±0.3‰ over 20 samples) (C). Calibration curve between N mass in acetanilide standards to the measured amplitude, used for quantification of the N in the sampled biomass (D). Symbols: Glutamic acid (USGS40) was marked by pink, Glycine (USGS64) by green, Caffeine (USGS62) by orange, and Acetanilide by grey.

*Preparation of lysis buffer.* Final concentrations of lysis buffer consisted of diethylpyrocarbonate water (DEPC Water) with dissolved 40 mM of

Ethylenediaminetetraacetic acid (EDTA), 50 mM of Trisaminomethane hydrochloric acid (Tris HCl, pH = 8.3) and 0.75 M of sucrose.

*NifH PCR*. The *nifH* genes were amplified using Dream Taq (EP0705, Thermo Scientific™) and primers (Hylabs, Israel) in a two-step polymerase chain reaction (PCR, Life Eco, Bioer Technology, China). In the first step, amplification was performed using the primers r: TTYTAYGGNAARGGNGG and f: ATRTTRTTNGCNGCRTA under the following conditions: 94 °C for 5 min, followed by 30 cycles of 94 °C for 1 min, 50 °C for 1 min, 72 °C for 1 min, and a final extension at 72 °C for 1 min. The second step utilized the primers r: ADNGCCATCATYTCNCC and f: TGYGAYCCNAARGCNGA under the following conditions: 94 °C for 5 min, followed by 30 cycles of 94 °C for 1 min, 57 °C for 1 min, 72 °C for 1 min, and a final extension at 72 °C for 1 min. PCR products were visualized using gel electrophoresis and validated with a positive control (*Vibrio natriegens*) and a negative control (double-distilled water, DDW).

### Results

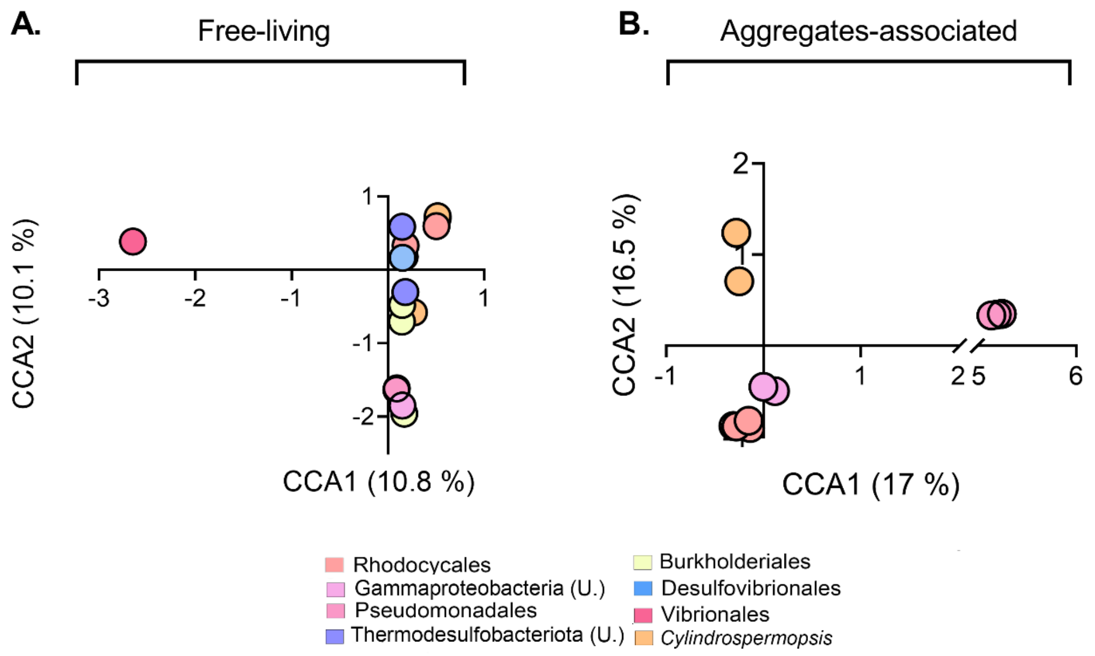


**Figure S2.** Beta-diversity was determined by a CCA model for free-living (A) and aggregate-associated (B) samples according to position of top 20 abundant orders of diazotrophs.

**Table S1.** Averages of primary production (PP) and bacterial production (BP) under light and dark treatments at 48 h from three sites of the upper Jordan River and Kinneret Lake.

| **Sampling location** | | **Upstream** | | **Midstream** | | **Downstream** | | **Lake** | |
|---|---|---|---|---|---|---|---|---|---|
| Season | | Summer | Winter | Summer | Winter | Summer | Winter | Summer | Winter |
| PP (μg C $L^{-1}$ $d^{-1}$) | Light | 1.4±0.2 | 1.6±0.8 | 4±3.1 | 7.8±3.1 | 34.6±20.4 | 3.3±1.2 | 85.1±30.7 | 85.7±41.5 |
| | Dark | 0.9±0.6 | 1.1±0.3 | 1.3±0.4 | 1±0.4 | 2±1.5 | 0.9±0.2 | 6.6±4.4 | 4±0.4 |
| BP (μg C $L^{-1}$ $d^{-1}$) | Light | 9±2 | 25.7±9.1 | 11.7±1.4 | 3.4±0.2 | 21.8±1.3 | 4.1±1.2 | 15.4±1.2 | 32.1±2.4 |

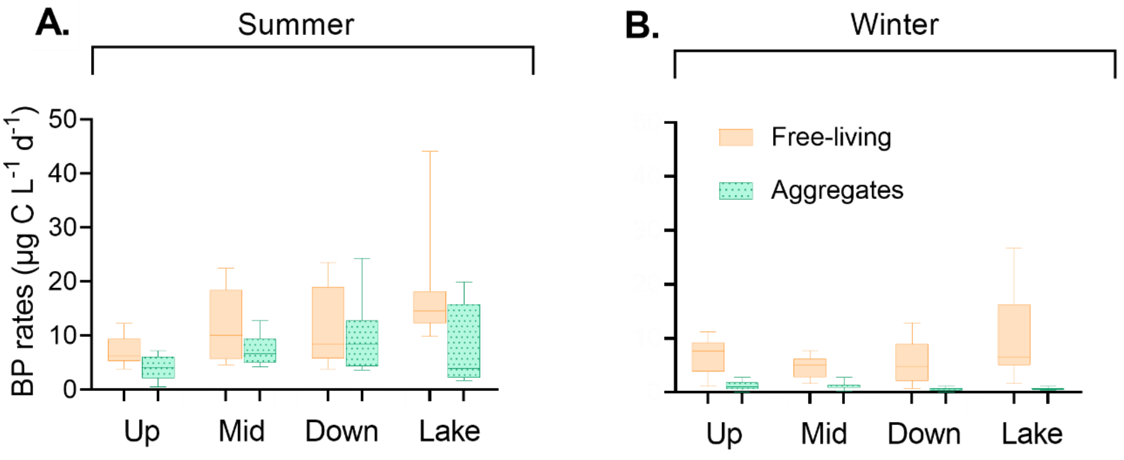


**Figure S3.** The spatial distribution of BP during Summer (A) and Winter (B) after 48 hours of incubation in the dark and divided into two size fractions free-living (orange, <12 µm) and aggregates associated (green, >12 µm)
